# Trans-biobank analysis with 676,000 individuals elucidates the association of polygenic risk scores of complex traits with human lifespan

**DOI:** 10.1101/856351

**Authors:** Saori Sakaue, Masahiro Kanai, Juha Karjalainen, Masato Akiyama, Mitja Kurki, Nana Matoba, Atsushi Takahashi, Makoto Hirata, Michiaki Kubo, Koichi Matsuda, Yoshinori Murakami, FinnGen, Mark J. Daly, Yoichiro Kamatani, Yukinori Okada

## Abstract

Human genetics seeks a way to improve human health on a global scale. Expectations are running high for polygenic risk scores (PRSs) to be translated into clinical practice to predict an inborn susceptibility to health risks. While risk stratification based on PRS is one way to promote population health, a strategy to utilize genetics to prioritize modifiable risk factors and biomarkers driving heath outcome is also warranted. To this end, here we utilized PRSs to comprehensively investigate the association of the genetic susceptibility to complex traits with human lifespan in collaboration with three worldwide biobanks (*n*_total_ = 675,898). First, we conducted genome-wide association studies for 45 quantitative clinical phenotypes, constructed the individual PRSs, and associated them with the age at death of 179,066 participants in BioBank Japan. The PRSs revealed that the genetic susceptibility of high systolic blood pressure (sBP) was strongly associated with a shorter lifespan (hazard ratio [HR] = 1.03, *P* = 1.4×10^-7^). Next, we sought to replicate these associations in individuals of European ancestry in UK Biobank (n = 361,194) and FinnGen (n = 135,638). Among the investigated traits, the individuals with higher blood pressure-related PRSs were trans-ethnically associated with a shorter lifespan (HR = 1.03, *P*_meta_ = 3.9×10^-13^ for sBP) and parental lifespan (HR = 1.06, *P*_UKBB_ = 2.0×10^-86^ for sBP). Further, our trans-biobank study identified additional complex traits associated with lifespan (e.g., obesity, height, serum lipids, and platelet counts). Of them, obesity-related traits showed strikingly heterogeneous effects on lifespan between Japanese and European populations (*P*_heterogeneity_ = 9.5×10^-8^ for body mass index). Through trans-ethnic biobank collaboration, we elucidated the novel value of the PRS study in genetics-driven prioritization of risk factors and biomarkers which can be medically intervened to improve population health.

## Introduction

Human disease risks can be explained by the combinations and interactions of inherited genetic susceptibility, acquired environmental exposures, and lifestyle factors^1^. One of the goals of medical research is to identify individuals at health risks both at the time of birth and later in life, and to provide them medical attention when necessary. Polygenic risk scores (PRSs) have successfully shown their predictive ability to idenitify those with a several-fold higher inherited risk of a given disease or condition^2^. Both an increase in statistical power and ethnic diversity in genetic studies—accelerated by nation-wide biobanks—have been instrumental in accurately predicting disease onset by PRSs^3–6^. Stratification of health risks based on PRSs would be one of the strategies to improve population health through targeted prevention. Nevertheless, the genetic risk itself cannot be modified. For many complex human traits, environmental exposure and lifestyle are also of great importance, such as cigarette smoking^7^ and dietary habits^8^. The accurate identification of risk factors that affect not only disease onset but also long-term health outcomes would contribute to population health, because these factors can be modified by medical intervention.

Observational studies have been attempting to identify monitorable risk factors and biomarkers that are correlated with the health outcomes (e.g., high low-density lipoprotein [LDL] cholesterol levels and the development of myocardial infarction). Nevertheless, the observational studies are inevitably laden with the pervasive issue of difficulty in inferring the cause-and-effect direction. A randomized controlled trial (RCT) is considered the gold standard to derive the effect of the exposure on the outcome free from unknown confounders^9^. In the above example, if a medical intervention to decrease the LDL cholesterol level leads to the decreased incidence of myocardial infarction at the population level, we could estimate that the high LDL cholesterol levels cause the development of myocardial infarction. The limitations of RCTs are, however, that they require a considerable amount of human and economic resources and are not always ethically feasible.

To address this, we here aimed to identify complex human traits affecting human lifespan, a health outcome of extreme importance and interest, by utilizing PRSs. The association of genetic susceptibility with lifespan would enable the prioritization of common risk factors and biomarkers, which could drive mortality in the current generation, among a variety of phenotypes that could be monitored in clinics. Furthermore, integration with deep-phenotype records and follow-up data in biobanks would enable us to pinpoint specific comorbidities and death causes that lead this association. Given the large genetic and environmental differences among populations, trans-ethnic comparison is also warranted. A collaboration with three trans-ethnic nation-wide biobanks collecting genotype, phenotype and survival data (*n*_total_ = 675,898) has enabled us to uncover the modifiable risk factors and monitorable biomarkers affecting human lifespan across the populations, on an unprecedented scale and without any clinical intervention.

## Results

### Study overview

An overview of our study design is presented in **Supplementary Figure 1**. We collaborated with three nation-wide biobanks (BioBank Japan [BBJ], UK Biobank [UKBB], and FinnGen) to elucidate clinical biomarkers affecting the lifespan of the current generation, across the different populations. The BioBank Japan cohort consisted of 200,000 participants mainly of Japanese ancestry, with clinical phenotype, biochemical measurement, lifestyle, and genotype data. The detailed information of this cohort is described elsewhere^10–12^ and in **Supplementary Table 1a**. Of them, 138,278 participants were followed up for their health record after an initial visit, including disease onset, survival outcome, and the cause of death if they died. The mean follow-up period was 7.44 years, and the number of deaths during the follow-up was 31,403. The UK Biobank project is a population-based prospective cohort consisting of approximately 500,000 people in the United Kingdom with deep phenotype and genotype data (summary in **Supplementary Table 1b**; see **URLs**). The biobank participants are linked to a death registry, which provides the age and cause of death when they die. In this study, we analyzed 10,483 deaths during a mean follow-up period of 6.97 years. FinnGen is a public-private partnership project combining genotype data from Finnish biobanks and digital health record data from Finnish health registries (see **URLs**). We analyzed 11,058 deaths in the national death registry among the 135,638 participants in this study.

We first sought to identify clinical biomarkers that were associated with lifespan in BioBank Japan and UK Biobank as an illustration of a conventional observational study. We then performed an association test of the PRSs (i.e., genetic susceptibility) of these biomarkers with lifespan in BioBank Japan, in order to elucidate the drivers, not the correlation, affecting human lifespan. We next performed replication studies of the association of the PRSs with lifespan in UK Biobank and FinnGen. We finally meta-analyzed these associations across the three cohorts.

### Association study of clinical biomarkers with human lifespan

First, in order to identify candidate clinical biomarkers correlated with human lifespan, we conducted an observational association study of these phenotypes with the lifespan on BioBank Japan. After the Bonferroni correction for multiple testing, 38 out of 45 clinical phenotypes showed a significant association with age at death (**Figure 1a**; Summary results are in **Supplementary Table 2**). The top traits associated with a shorter lifespan were low albumin, high γ-glutamyl transpeptidase, and increased height. The effect of a one standard deviation (SD) increase in each trait on mortality resulted in a hazard ratio (HR) of 0.80 [0.79– 0.81], 1.16 [1.15–1.17], and 1.30 [1.27–1.32] (*P* = 3.3 × 10^−287^, 1.1 × 10^−224^, and 8.3 × 10^−186^), respectively. These results were consistent with the previous epidemiological studies in other cohorts^13–16^.

**Figure 1.**
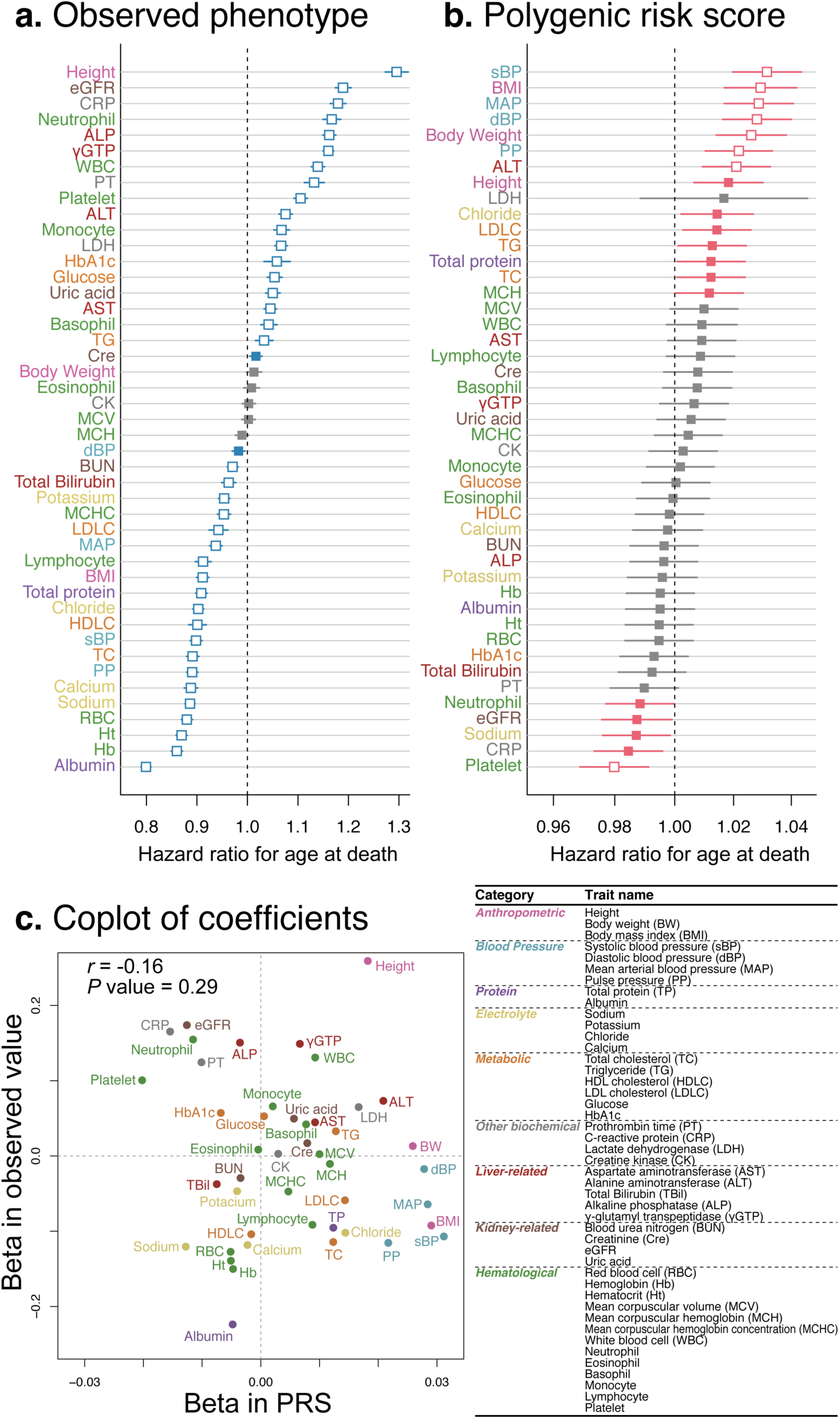
The hazard ratios for the age at death, according to clinical phenotypes and according to the PRSs and their correlations in BioBank Japan. Shown are the adjusted HRs from Cox proportional-hazard models for lifespan, according to clinical phenotypes **(a)** and according to the PRSs for the clinical phenotypes **(b)** in BioBank Japan. The boxes indicate the point estimates, and the horizontal bars indicate the 95% confidence interval. Boxes in blue **(a)** or red **(b)** indicate the nominal significance (*P* < 0.05), and the white-out boxes indicate the statistical significance after correcting for multiple testing by the Bonferroni method. **(c)** Co-plot of the coefficients from the Cox proportional-hazard models for lifespan according to the PRS (x-axis) and those according to clinical phenotypes (y-axis).

To investigate how the association of clinical biomarkers with human lifespan is shared across different populations, we next performed the same observational study in UK Biobank using the 20 clinical phenotypes that were recorded in both UK Biobank and in BioBank Japan (**Supplementary Figure 2**). We again observed significant associations of the quantitative traits with lifespan in 17 out of 20 traits. Of note, 14 among the 15 traits with significant association in BioBank Japan showed directionally concordant associations with lifespan in UK Biobank. The only trait that showed directionally discordant association was body mass index (BMI). While a lower BMI was significantly associated with a shorter lifespan in Biobank Japan, a higher BMI showed significant association with a shorter lifespan in UK Biobank. This discordant result could be attributed to differences in the participation criteria (i.e. hospital-based recruitment in BioBank Japan and healthy volunteers in UK Biobank) and differences in the health burden of obesity across populations, which warrants further replication studies in different cohorts.

A weakness of the epidemiological associations was, however, that it was difficult to conclude whether the variations in clinical measurements had caused the variations in lifespan, or they were just correlations. For example, a decreased albumin level was associated with a shorter lifespan, but this did not mean that low albumin caused high mortality. Rather, the decline in general health and nutritional status, which led to high mortality, might have resulted in low albumin levels.

### Association study of PRSs of complex traits with human lifespan in BioBank Japan

Next, in order to prioritize the clinical traits affecting human lifespan, we utilized genetic information. PRS is supposed to simulate the genetic predisposition towards the investigated trait^1^. Thus, the association of the PRS of the investigated trait with lifespan can be considered as less susceptible to the confounding factors such as a decline in general health^17, 18^. PRSs should be constructed from the genetic studies of the same population^5, 19^, and BioBank Japan has been the largest study of East Asian populations to date. Conventionally, when independent large-scale GWAS statistics with matched population and a sufficient sample size are not available for constructing PRSs, a strategy to split the study cohort into two groups (i.e., discovery group to conduct GWASs and a validation cohort to derive the PRSs) has been used. This strategy compromises accurate estimates in GWAS statistics using maximum samples or lowers the statistical power in PRS validations, depending on how the cohort is split. To address this, we adopted a 10-fold leave-one-group-out (LOGO) meta-analysis approach in the derivation of PRSs in order to validate the PRSs in participants independent from GWAS while retaining as much sample size and statistical power as possible. Briefly, we first conducted GWASs on 45 clinical phenotypes by randomly splitting the whole cohort into ten sub-groups (**Supplementary Table 3** shows the detailed phenotype information used in GWASs). Then, we meta-analyzed nine GWASs (**Supplementary Table 4** for the GWAS summaries), constructed PRSs from the meta-analyzed statistics by using a clumping and thresholding method, and performed survival analyses to investigate the association of the derived PRS with individual lifespan (age at death) in the one withheld sub-group. We repeated this analysis ten times and further meta-analyzed the statistics of survival analyses in the 10 sub-groups (**Figure 2** and **Methods** for the study design). Thus, we were able to maintain the sample size in GWASs at nine-tenths of the whole cohort and at the same time, validate the derived PRSs using all of the individuals in the cohort.

**Figure 2.**
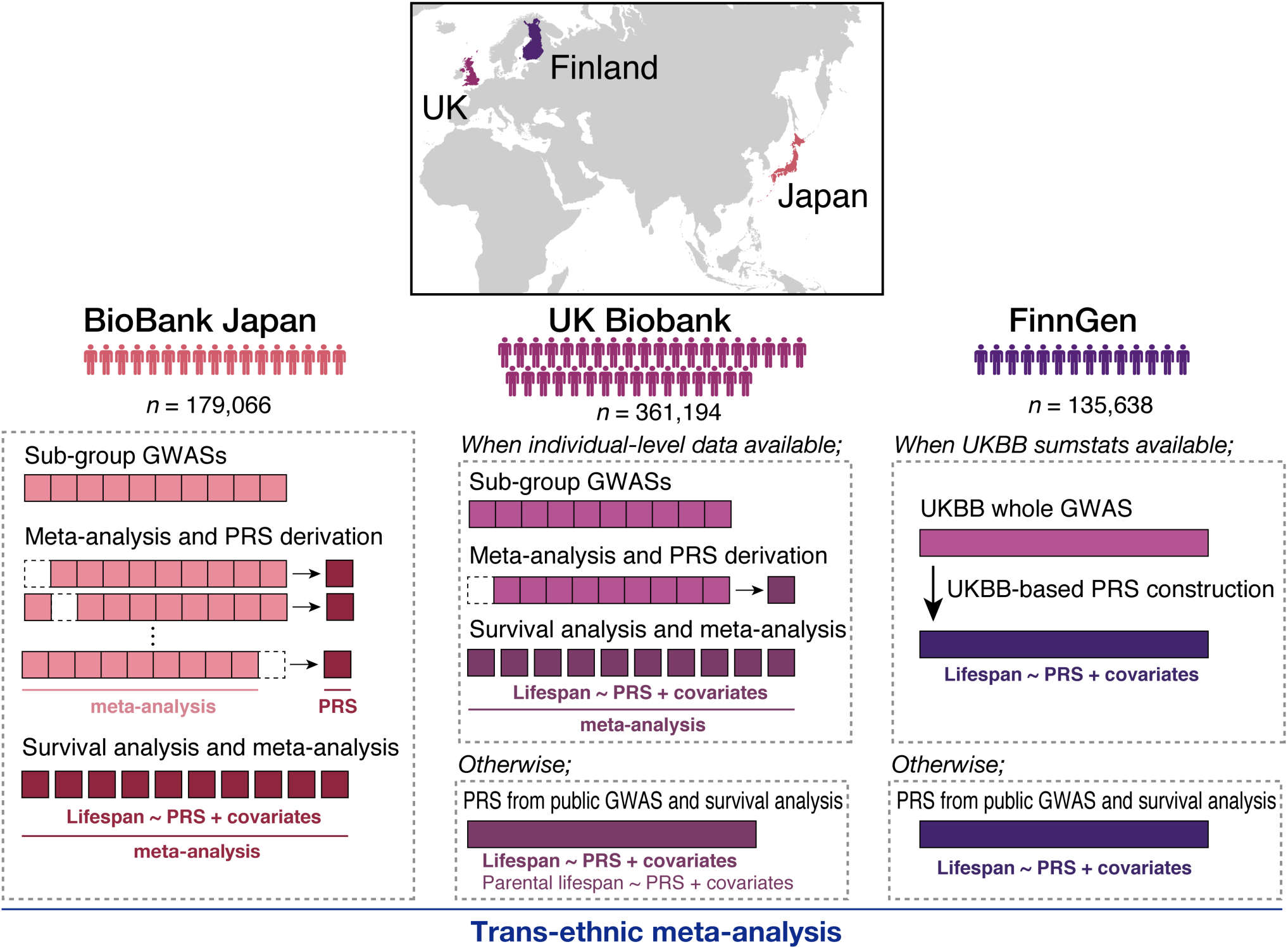
Overview of PRS-lifespan association study in collaboration with three nation-wide biobanks. In BioBank Japan, we first randomly split the entire cohort into 10 sub-groups and performed genome-wide association studies (GWASs) on 45 quantitative traits. We then performed a 10-fold leave-one-group-out (LOGO) meta-analysis, derived the PRSs in one remaining sub-group, and associated them with lifespan. We meta-analyzed the statistics of the lifespan association obtained from the ten sub-groups. In UK Biobank, when individual-level phenotype data is available, we adopted the LOGO approach. Otherwise, we derived the PRSs from public large-scale GWAS statistics, and associated the PRSs with lifespan in the cohort. As a secondary analysis, we also associated the PRS with parental lifespan in UK Biobank. In FinnGen, we derived the PRSs from UK Biobank GWAS summary statistics or public large-scale GWAS statistics, and associated the PRS with lifespan in the cohort. Finally, we performed trans-ethnic meta-analysis.

Among the investigated clinical phenotypes, higher PRSs of blood pressure-related traits (systolic blood pressure [sBP], diastolic blood pressure [dBP], and mean arterial pressure [MAP]) were significantly associated with a shorter lifespan (**Figure 1b;** summary results shown in **Supplementary Table 2**). In the case of sBP, whose PRS showed the strongest association with the age at death (HR of per SD increase in PRS on mortality = 1.03 [1.02– 1.04], *P* = 1.4×10^-7^), individuals with the highest sBP PRS (in the top quintile) had indeed a 1.46-fold higher risk of being hypertensive (sBP > 130 mmHg or dBP > 80 mmHg) or being treated with anti-hypertensive medications when compared with those with the lowest PRS (in the bottom quintile; *P* = 1.4×10^-84^). A comparison between the standardized survival curves according to the observed phenotype and those according to the PRS of the phenotype is highlighted in **Figure 3**. Those with the highest PRS (in the top quintile) and thus with the genetic predisposition to cause increased blood pressure were significantly associated with an increased risk of standardized mortality than those with the lowest PRS (the standardized 10-year mortality rate was 0.210 and 0.217, respectively, **Figure 3b**, top). On the other hand, the measured blood pressure value showed U-shaped associations with lifespan, with those with the lowest and the highest sBP both harboring an increased risk of mortality (**Figure 3a** top and **Supplementary Figure 3**). By utilizing the genetic data, we disentangled the dose-dependent association of the genetic risk of high blood pressure with a short lifespan, while the observed association of the lowest-range blood pressure with a short lifespan might have been confounded by the consequence (i.e. decline in general health caused low blood pressure^20^). This contrasts with the case of albumin. Although the measured low albumin level showed the strongest association with a short lifespan (**Figure 3a**, bottom), the PRS of the albumin did not show any association with the age at death (HR = 0.99 [0.98–1.00], *P* = 0.40, **Figure 3b**, bottom). Overall, there was no significant correlation of the effect size and directions between the association of clinical phenotypes on lifespan and the association of PRSs of clinical phenotypes on lifespan (*r* = −0.16, *P* = 0.29; **Figure 1c**), which was not confounded by the variance explained by PRSs in each trait (shown in **Supplementary Table 5**). To summarize, the PRSs have provided novel and distinct insights into prioritizing critical factors affecting human lifespan from the observational studies.

**Figure 3.**
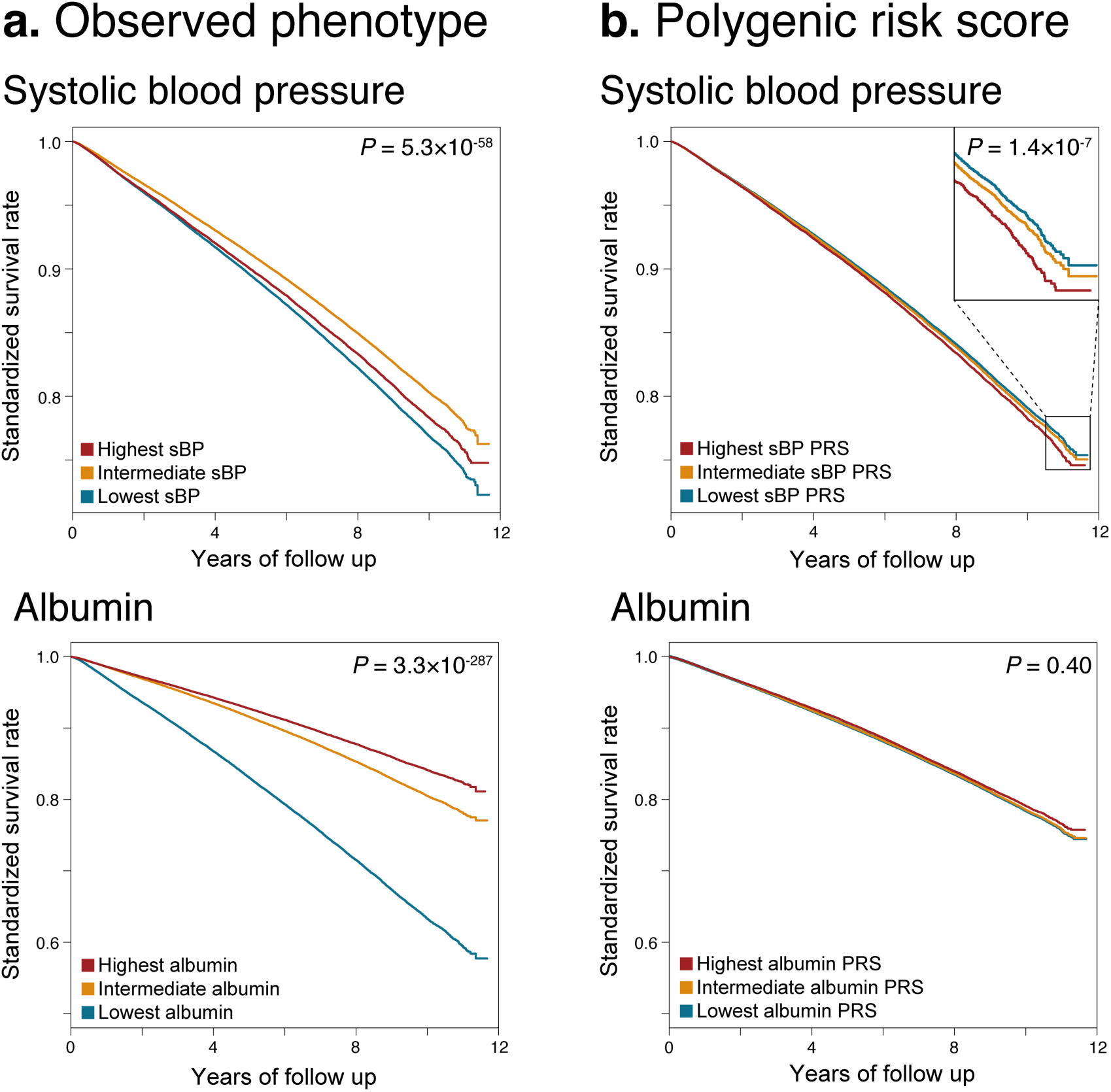
Standardized survival rate, according to systolic blood pressure (sBP) and albumin, and PRS status of both traits in BioBank Japan. In each box, the standardized and adjusted survival curves according to three bins (lowest, first quintile; intermediate, 2–4 quintiles; and highest, fifth quintile) of the investigated trait or the PRS of the investigated trait are illustrated by analyzing mortality data in BioBank Japan. The standardization was performed using the mean of all the covariates. **(a)** Survival curves according to measured sBP value (top) and according to sBP PRS (bottom). **(b)** Survival curves according to measured serum albumin level (top) and according to albumin PRS.

In addition to the overall survival outcome, we also tested the cause-specific mortality that drives the association with the sBP PRS, by leveraging the detailed follow-up data in BioBank Japan. Among the four most frequent causes of death in Japan^11^, a high sBP PRS was significantly associated with death from cardiovascular diseases (I01–I02, I05–I09, I20–I25, I27 and I30–I52 [HR = 1.04 (1.01–1.08), *P* = 0.0064]) and nominally associated with death from cerebrovascular diseases (I60–69 [HR = 1.05 (1.01–1.10), *P* = 0.024]), as categorized by the International Classification of Diseases 10. We next performed comorbidity-stratified analysis in the association of sBP PRS with lifespan. We found that individuals with a past medical history of type 2 diabetes, cerebral infarction, or dyslipidemia strongly drove the association of sBP PRS with lifespan in Japanese individuals (HR = 1.05 [1.03–1.07), 1.06 [1.03–1.09), 1.05 [1.02–1.08), and *P* = 2.6×10^-5^, 1.9×10^-4^, 4.0×10^-3^, respectively). These results recapitulated the epidemiological knowledge that high blood pressure is one of the strongest risk factors of mortality among patients with cardiovascular^21^, cerebrovascular ^22, 23^, and metabolic diseases^24^. It has been previously reported that healthy-aging individuals had low genetic risk of coronary artery disease^25^, which is in line with our findings.

### Trans-ethnic association study of PRSs of complex traits with human lifespan

Next, we sought to replicate these associations in individuals of European ancestry using the individual-level data of UK Biobank (*n* = 361,194) and FinnGen (*n* = 135,638). We first constructed the PRSs by adopting the 10-fold LOGO meta-analysis approach when the individual-level phenotype was available (20 out of 33 traits in UK Biobank). Otherwise, we derived the PRSs by using independent publicly available large-scale GWAS summary statistics of European ancestry (13 out of 33 traits in UK Biobank and all the 33 traits in FinnGen) with a linkage disequilibrium (LD) reference of European individuals (**Figure 2** and **Methods** for the study design and **Supplementary Table 6** for public GWAS information). In this way, we could calculate the individual PRSs of 33 quantitative traits among the 45 investigated traits in BioBank Japan (**Supplementary Table 7** and **8** for phenotype and internal GWAS summary). We then associated the derived PRSs with lifespan in UK Biobank and FinnGen, and finally performed a trans-ethnic meta-analysis across the three cohorts (Summary results are shown in **Supplementary Table 9**). In UK Biobank and FinnGen, we successfully replicated the directional consistency of the association of a genetically increased risk of sBP with a shorter lifespan (HR = 1.02 [1.00–1.04], *P* = 0.083 in UK Biobank [**Figure 4b**] and HR = 1.03 [1.01–1.05], *P* = 0.0031 in FinnGen [**Figure 4c**]). A fixed-effect meta-analysis revealed a trans-ethnically robust association of higher PRSs of sBP with a shorter lifespan (HR = 1.03 [1.02–1.04], *P* = 3.9×10^-13^; **Figure 4d**). To further validate this finding, we also performed a secondary analysis using parental lifespan data in UK Biobank, which offered a much larger statistical power (see **Methods** for the detailed analysis method). The secondary analysis revealed that a genetically increased risk of sBP was also associated with a shorter parental lifespan (HR = 1.06 [1.06–1.07], *P* =2.0×10^-86^).

**Figure 4.**
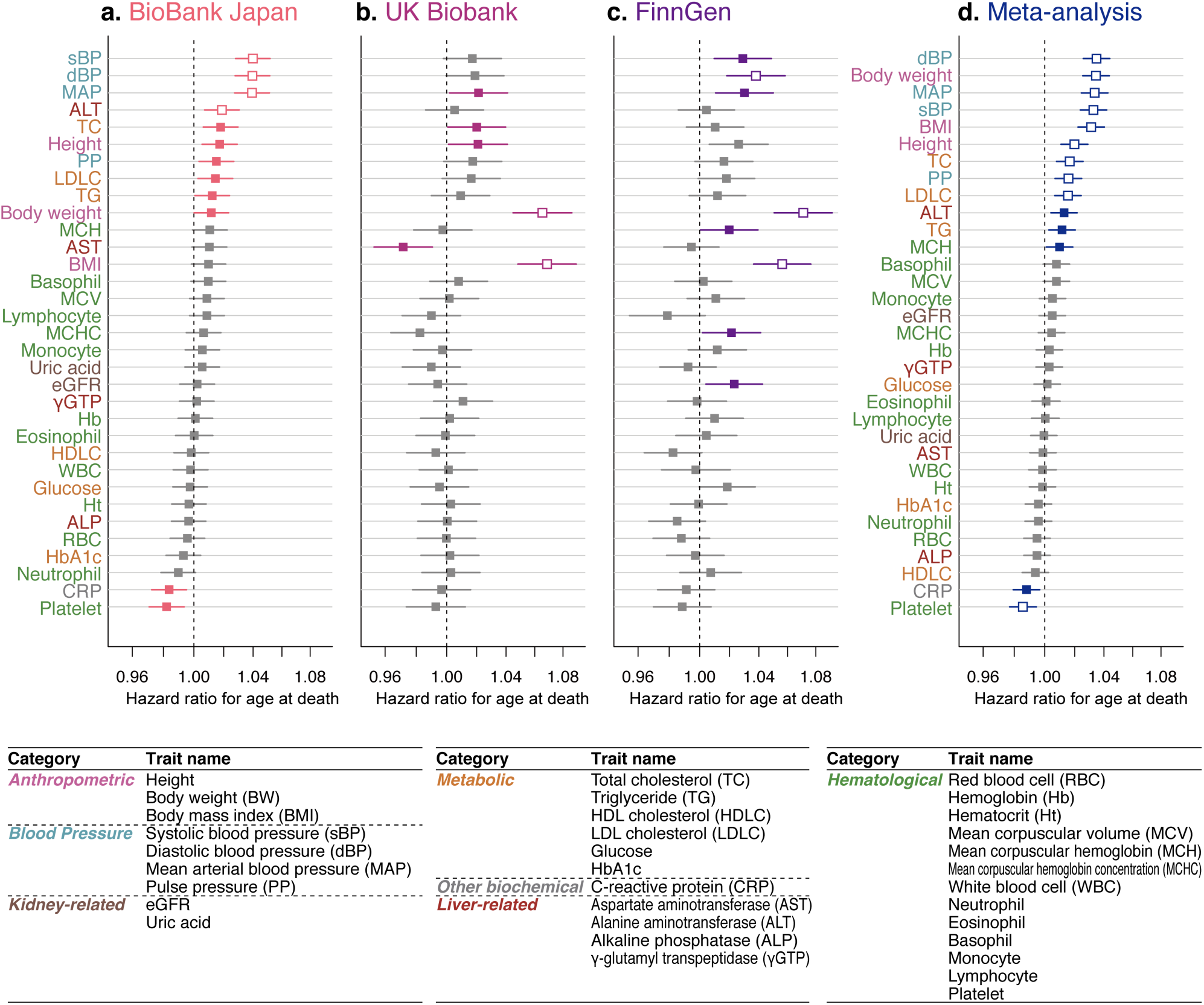
Trans-ethnic association study of PRS with lifespan. Shown are the adjusted HRs from Cox proportional-hazard models for lifespan, according to the PRS of the clinical phenotypes in **(a)** BioBank Japan, **(b)** UK Biobank and **(c)** FinnGen. The threshold of significance for the derivation of PRS was set *P* = 1.0×10^-6^. We further performed a trans-ethnic fixed-effect meta-analysis of the association results from the three cohorts **(d)** by the inverse-variance method. The boxes indicate the point estimates, and the horizontal bars indicate the 95% confidence interval. Boxes in colors indicate the nominal significance (*P* < 0.05) and the white-out boxes indicate the statistical significance after the Bonferroni correction for multiple testing (*P* < 1.5×10^-3^).

Interestingly, the high PRSs of BMI and body weight (BW) were most significantly associated with short lifespan in UK Biobank and FinnGen (BMI: HR = 1.07 [1.05-1.09] and 1.06 [1.04-1.08], *P* = 1.7×10^-11^ and 1.5×10^-8^, respectively), while they showed much smaller effect sizes and less significant associations in BioBank Japan (BMI: HR = 1.01 [1.00-1.02], *P* = 0.094). We noted that a strong effect of obesity on lifespan was consistent between the two of the European cohorts, UK Biobank and FinnGen, which would suggest the robustness of the result against the methods used for the calculation of PRSs (i.e., LOGO in UK Biobank and usage of independent GWAS summary statistics in FinnGen). Among all the investigated traits, the random effect meta-analyses only revealed a significant heterogeneity in association for BMI and BW (*P*_heterogeneity_ = 9.5×10^-8^ [BMI] and 1.5×10^-8^ [BW]). We did not observe apparent differences in the heritability and variance explained by the PRSs of BMI or BW between in BioBank Japan and UK Biobank (**Supplementary Table 4, 5, 8 and 10**). Thus, we considered that the reasons for this trans-ethnic heterogeneity was not attributed to the differences in GWASs utilized for the derivations of PRSs. The observed trait mean and SD were larger in the European cohorts before normalization (the mean for BMI was 23.3, 27.4, and 27.2, and the SD was 3.7, 4.8, and 4.1 in BioBank Japan, UK Biobank and FinnGen, respectively), and this was also the case in World Health Organization (WHO) data from 2016 (BMI: 22.8 [22.5–23.2] in Japan, 27.5 [27.2–27.8] in UK, and 26.6 [26.1–27.1] in Finland; see **URLs**). Obesity-related cardiovascular deaths are significantly prevalent among Europeans, and the epidemiological data revealed that the mortality rate of individuals in the in high BMI range was higher in Europeans than in East Asians^26^. The heterogeneity in the association of BMI or BW PRSs on lifespan between Japanese and Europeans might reflect the differences in the strength of the effect of obesity on mortality, on which further trans-ethnic studies should be warranted.

To determine what is driving the association of BMI PRS on lifespan (i.e. mortality) in Europeans, we additionally investigated the cause-specific mortality and comorbidity information recorded in UK Biobank. When we tested the association of BMI PRS with the cause-specific mortality in UK Biobank, the BMI PRS was most strongly associated with cerebrovascular death (HR = 1.12 [1.08–1.17], P = 3.1×10^-8^). When we stratified individuals based on the comorbid conditions (i.e., common disease affection status), we found that the association of BMI PRS with lifespan was strongest among those with unstable angina (HR = 1.17 [1.05–1.30], P = 3.1×10^-3^). These analyses successfully pinpointed the target individuals who would be expected to benefit most from the modification of obesity.

There were several additional traits where the PRSs showed significant associations with lifespan in trans-ethnic meta-analysis after the Bonferroni correction for multiple testing (*P*_meta_ < 1.5×10^-3^; i.e., lipid-related traits, height, and platelet count). The genetic burden of increased lipid-related traits (i.e. total cholesterol and LDL cholesterol) was associated with a shorter lifespan, which was concordant with the observational studies reporting the causal roles of cholesterol in worse health outcomes^27^. Height has been indicated as a risk factor for various cancers and linked with cancer-related mortality in both Europeans and Asians^15, 16^. A lower platelet count was also reported as associated with an increased mortality in Europeans^28^.

To test whether there existed differences in the effect sizes of PRS on lifespan between males and females, we performed a sex-stratified association study of PRSs with lifespan across the investigated traits and across the three cohorts. While we did not find any significant differences between sexes within each of the three cohorts (**Supplementary Figure 4a–c**), the sex-stratified trans-ethnic meta-analysis revealed that the effect of high dBP PRS on a short lifespan, which was the largest among 33 traits in primary meta-analysis, was significantly larger in males than in females (HR_male_ = 1.05 [1.04-1.06], HR_female_= 1.02 [1.00-1.03], *P*_heterogeneity_ = 0.0013; **Supplementary Figure 4d**). This observation was in line with previous epidemiological studies showing that the excess mortality caused by hypertension was higher for men than for women in the Japanese population^29^, and that women with hypertension not complicated by left ventricular hypertrophy had lower risk of clinical major cardiovascular events than men in Europeans^30^.

Finally, in order to validate our findings, we conducted a trans-ethnic Mendelian randomization (MR) study of the 33 traits on which we had performed the trans-ethnic PRS-lifespan association study (see **Methods**). Two-sample MR with the inverse-variance weighted method revealed the following; (i) the significant causal effect of sBP and mean arterial blood pressure (MAP) on lifespan in BioBank Japan; (ii) the significant causal effect of BMI and BW on lifespan in UK Biobank and FinnGen; and (iii) that trans-ethnic meta-analysis further strengthened their significance (i.e. BMI, BW, sBP, and MAP; *β*_causal_ = 0.17, 0.17, 0.15, 0.15; and *P*_meta_ = 1.6×10^-11^, 9.6×10^-11^, 1.6×10^-4^, 8.2×10^-4^ respectively; summary results shown in **Supplementary Figure 5**). While both methods (PRS and MR study) have their own limitations, such as pleiotropy and assumptions on instrumental variables^31, 32^, we consider that the consistent result from these two methods would complement each other and further support the robustness of our findings in identifying the driver biomarkers of human lifespan.

To summarize, these results collectively suggest the utility of PRSs in genetics-driven identification of both known and novel drivers for longevity, and could potentially pinpoint a group of individuals who could most likely benefit from the intervention.

### No evidence of an interaction effect of PRSs of complex traits and lifestyle factors on lifespan

Motivated by the identification of biomarkers genetically affecting lifespan, we finally investigated whether there existed any interaction between the PRS of these biomarkers and various lifestyles. As blood pressure PRSs were most strongly associated with lifespan in the Japanese population, we tested the interaction effect between sBP PRS and lifestyle on lifespan in BioBank Japan (**Supplementary Table 11**). While various lifestyle factors had a strong impact on lifespan (**Supplementary Table 12**), none of them showed significantly heterogeneous effects on survival according to the sBP PRS status. For example, the beneficial effect of smoking cessation on survival was not significantly different among those with the highest risk of increased blood pressure (Δ10-year mortality = −0.050) or those with the lowest risk of increased blood pressure (Δ10-year mortality = −0.049, interaction *P* = 0.63) inBioBank Japan. In Europeans, as we found the strongest association between the obesity PRS and lifespan, we investigated the interaction between BMI PRS and lifestyle in UK Biobank. Again, no significant interaction effect on lifespan was observed (*P*_interaction_ > 0.05).

Taken together, even people with the high genetic burden of increased blood pressure or obesity could benefit from the modifiable lifestyles such as abstinence from smoking and regular exercise, which could lead to a better survival.

## Discussion

Harnessing a global effort to expand genetic studies in both sample size and the scope of phenotypes, with the additional notion of the importance of population diversity^5^, PRS is expected to identify individuals with inborn health risks in clinics. While early detection and appropriate health communication should contribute to the improvement of health care^33^, the inherited genetic risks of disease onset cannot be modified.

We here showed the novel value of PRS study to identify the monitorable phenotypes that genetically affect health outcomes. Our approach has the potential to contribute to the improvement of healthcare because the identified factors can be modified by medical intervention. We showed a global burden of increased blood pressure and obesity as drivers of mortality from genetics. Our study also revealed that those with a genetic burden to cause high blood pressure or obesity could benefit from healthy lifestyles to the same degree as those without. If those with high-risk alleles are to be notified about their own risks, the early lifestyle modification and medical attention should prevent their premature death. Of note, the magnitude of the effect size in which the PRS of the trait was associated with lifespan was relatively small. However, the magnitude of effect size in which the trait itself (e.g., blood pressure or obesity) affects lifespan, or in which the modification of the trait (e.g., proper blood pressure management or healthy diet) would improve health outcomes, would be expected to be larger in terms of population health.

In order to improve population health, we need to decide on how to prioritize the numerous health issues. The observational studies could partly address this point, but the biggest challenge has been that we cannot infer the cause-and-effect direction. While RCTs have been the gold standard to provide robust evidence of the effect of risk factors on health outcomes, they are not always feasible because conducting RCTs (i.e. recruitment, random allocation, treatment, and follow-up etc.) takes a huge amount of resources, which hampers the application to diverse phenotypes. Our approach, which leverages genetic and phenotypic information already existing in biobanks, would have the potential to support the clinical evidence, or to identify candidate risk factors to bring into RCTs. We also note that in-depth analyses, such as those leveraging cause-specific mortality and comorbidity data, could pinpoint target individuals who could most likely benefit from medical attention and intervention. These insights would also be useful in designing efficient RCTs or providing individualized medical evidence.

Notably, the genetics-driven identification of critical factors for health outcomes was made possible by trans-ethnic, large-scale, and deep-phenotyped biobanks. The trans-biobank collaboration provided (i) a large sample size, which was critical in analyzing mortality data, (ii) the opportunity for replication, which made our findings robust to cohort-specific confounders, (iii) a trans-ethnic comparison as in the example of obesity, (iv) the validation of our methodology (i.e., we confirmed the coherent result between LOGO and independent GWAS), and (v) the integration of cohort-specific data, such as parental lifespan data in UK Biobank. Nation-wide biobanks, such as those in this study, are prospectively collecting deep phenotype and health outcomes of genotyped individuals, and our proof-of-concept approach would be expected to discover the actionable traits driving health outcomes on a global scale if further applied to diverse and larger populations.

This study has potential limitations. First, as BioBank Japan is a hospital-based cohort, it does not represent the Japanese population as a whole. However, since we performed the survival analyses with an adjustment for the disease status and principal components followed by sensitivity analyses, our main result was not confounded by the proportion of patients with a specific disease group (**Supplementary Figure 6**) or population stratification. Of note, UK Biobank generally enrolls healthy-volunteers^34^. The directional concordance of statistics in BioBank Japan with those in UK Biobank should further support the robustness of the results and mitigates the concern on potential biases due to the differences in genetic structure and environmental interactions. Second, it is unclear whether the PRSs of the traits that showed less significant results in our study were not associated with lifespan because there is truly no relationship, because the PRS did not sufficiently explain the variance of the investigated phenotype, or because there was a strong effect of rare variants, which were not captured in our study. Third, the polygenic effect of the variants constituting the PRSs which also partially affect other traits (pleiotropy), might have coexisted with the association of the PRS of a specific trait with lifespan. Further integration with novel statistical methods to handle and disentangle pleiotropy and desirably RCTs, if feasible, are warranted to obtain clearer insights into the true effect of the complex trait on human lifespan. Fourth, there is currently no consensus on how to optimize and harmonize the *P* value threshold in calculating PRSs across different cohorts. Our strategy was to set a fixed *P* value threshold of 1×10^-6^ in the trans-ethnic meta-analysis, because we could not obtain trait-specific best *P* values for every trait which should be optimized to maximize the variance explained by using individual-level phenotype data. We confirmed that association statistics (i.e. coefficients) from the fixed threshold of 1×10^-6^ was fairly concordant with those from best *P* values (Pearson’s *r* = 0.85 and *P* = 2.5×10^-13^ in BioBank Japan, and *r* = 0.93 and *P* = 1.3×10^-9^ in UK Biobank). Nevertheless, we consider that further implementation of the methodology for optimally harmonizing PRSs across different cohorts is still warranted. Fifth, it is possible that spouse-pairs in biobanks might have caused a subtle bias the GWAS and PRSs-lifespan association if assortative mating exists^35^. Sixth, although we exhaustively checked the cohort-level overlap across biobanks and previous GWASs used in this study, we could not completely exclude the possibility of individual-level overlap, which would be technically difficult to detect as a general point in large-scale genetic studies. Last, the statistical power in the association study with lifespan was limited, partly due to a relatively short follow-up period. This was particularly the case in UK Biobank, which is a recently launched population-based cohort, and only a small number of people have died during the follow-up period. We complemented this point by utilizing parental lifespan data in UK Biobank as a secondary analysis. Since the participants of the biobanks in this study are ongoingly followed-up, the larger number of mortality records in the future would provide us with an opportunity to further validate the robustness of our results.

In conclusion, through trans-ethnic biobank collaboration, we demonstrated that blood pressure and obesity were genetically associated with the lifespan of the current generation on a global scale. A comparison across different populations and the integration with deep phenotype data further pinpointed a group of individuals who would be expected to benefit most from the intervention of these traits. With global biobanks’ ongoing efforts—enrolling individuals from diverse background and collecting granular phenotype along with health outcomes—we have shown a potential application of genetics to improve population health by providing information of common and modifiable risk factors driving our health outcomes.

## Acknowledgments

We sincerely thank all the participants of BioBank Japan, UK Biobank, and FinnGen. We thank Dr. Aarno Palotie for his kind support for the data analysis of FinnGen, Drs. Benjamin M. Neale and Nikolas Baya for sharing and discussing their idea on LOGO, and Dr. Alicia R. Martin for the PRS analysis on UK Biobank.

## Author Contributions

S.S., Y.K., and Y.O. conceived the study. M.H., M. Kubo, K.M., Y.M. collected and managed the BioBank Japan samples. S.S., M. Kanai, M.A., N.M., A.T., M. Kubo., Y.K. and Y.O. performed data cleaning and statistical analysis on Biobank Japan. M. Kanai performed statistical analysis on UK Biobank. J.K., M. Kurki and M. Kanai performed data cleaning and statistical analysis on FinnGen. M. J. D. contributed to the overall study design and the FinnGen analysis. Y.O. supervised the study. S.S., M. Kanai, J.K., Y.K., and Y.O wrote the manuscript.

## Competing Financial Interests

The authors declare that no conflicts of interest exist.

## Methods

### Study Populations, genotyping and imputation

#### BioBank Japan

Clinical information and genotype data were obtained from BioBank Japan (BBJ) project^10, 12^, which is a prospective biobank that collaboratively collected DNA and serum samples from 12 medical institutions in Japan and recruited approximately 200,000 participants, mainly of Japanese ancestry, with the diagnosis of at least one of 47 diseases. All the participants provided written informed consent approved from ethics committees of RIKEN Center for Integrative Medical Sciences, and the Institute of Medical Sciences, the University of Tokyo. Detailed participant information is summarized in **Supplementary Table 1a**.

We genotyped participants with the Illumina HumanOmniExpressExome BeadChip or a combination of the Illumina HumanOmniExpress and HumanExome BeadChips. The quality control (QC) of participants and genotypes was described elsewhere^36^. In this project, we analyzed 179,066 participants of Japanese ancestry as determined by the principal component analysis (PCA)-based sample selection criteria. The genotype data was further imputed with 1000 Genomes Project Phase 3 version 5 genotype (*n* = 2,504) and Japanese whole-genome sequencing data (*n* = 1,037)^36^ using Minimac3 software. After the imputation, we excluded variants with an imputation quality of Rsq < 0.7 or those with a minor allele frequency (MAF) < 1%.

#### UK Biobank

The UK Biobank project is a population-based prospective cohort that recruited approximately 500,000 people aged between 40–69 years from 2006 to 2010 from across the United Kingdom (summary in **Supplementary Table 1b**; see **URLs**). Deep phenotype data, such as electronic medical records, lifestyle indicators and bioassays, and genotype data were available for most of the participants. The genotyping was performed using either the Applied Biosystems UK BiLEVE Axiom Array or the Applied Biosystems UK Biobank Axiom Array. The genotypes were further imputed using a combination of the Haplotype Reference Consortium, UK10K, and 1000 Genomes Phase 3 reference panels by IMPUTE4 software. The detailed characteristics of the cohort were previously extensively described^34^. In this project, we analyzed 361,194 individuals of white British genetic ancestry as determined by the PCA-based sample selection criteria (see **URLs**). We excluded the variants with (i) INFO score ≤ 0.8, (ii) MAF ≤ 0.0001 (except for missense and protein-truncating variants annotated by VEP^37^, which were excluded if MAF ≤ 1 × 10^-6^), and (iii) HWE *P* ≤ 1 × 10^-10^. All of the analyses were conducted via application 31063.

#### FinnGen

FinnGen is a public-private partnership project combining genotype data from Finnish biobanks and digital health record data from Finnish health registries (see **URLs**). Six regional and three country-wide Finnish biobanks participate in FinnGen. Additionally, data from previously established population and disease-based cohorts are utilized. Participants’ health outcomes are followed up by linking to the national health registries (1969–2016), which collect information from birth to death. We used the genotype and phenotype data of 135,638 participants in this study, excluding population outliers via PCA (summary in **Supplementary Table 1c**). These individuals were genotyped with the FinnGen1 ThermoFisher array and previous cohorts were genotyped with various genotyping arrays. The genotype data was imputed using whole genome sequencing data from 3,775 Finnish individuals by beagle4.1 software (see **URLs**)^38^. After the imputation, we excluded variants with an imputation INFO score < 0.8 or MAF < 0.0001.

### Survival analysis of clinical phenotypes

We used Cox proportional hazard models to test the association of clinical phenotypes with lifespan (age at death) in BioBank Japan as described elsewhere^39^. In order to obtain and compare the HRs for the all-cause mortality across the traits, we scaled each trait to have zero mean and unit variance by Z-score transformation. The primary analyses included adjustment for sex, the 47-disease status and the top 20 principal components, which were supposed to account for possible confounders and population stratification. Additional summaries of clinical phenotypes and the number of samples without missing values are described in **Supplementary Table 3**. We next performed the same survival analyses in 20 clinical phenotypes where individual-level phenotype data was available in UK Biobank (**Supplementary Table 7**). We used Cox proportional-hazard models to test the association of these clinical phenotypes with lifespan (age at death) with an adjustment for sex and the top 20 principal components as covariates.

### Genome-wide association studies

#### BioBank Japan

In order to derive population-specific PRSs of BioBank Japan, we first split the cohort into ten sub-groups. We then conducted GWASs for 45 quantitative traits within each of the ten sub-groups. We performed the linear regression assuming the additive effect of the imputed dosage of each variant by PLINK^40^. For individuals taking anti-hypertensive medications, we added 15 mmHg to their sBP and 10 mmHg to their dBP and derived their MAP and pulse pressure (PP) using the adjusted sBP and dBP. We also added smoking status as a covariate for blood pressure-related traits. Other trait-specific covariates, adjustment for medications, and sample exclusion criteria are described in **Supplementary Table 13** and elsewhere^41^. We next meta-analyzed the statistics from nine sub-groups by the inverse-variance method assuming the fixed-effect ten times, with keeping one sub-group away from the meta-analysis for PRS derivation and validation each time (a 10-fold leave-one-group-out [LOGO] meta-analysis approach). Before performing LOGO, we excluded genetically related individuals from the cohort, based on PI_HAT > 0.125, as calculated by PLINK software. We note that we adopted this strategy to obtain precise estimates of the HR, not to maximize *R^2^* value, which will be maximized when we have the largest GWAS samples. We applied LD Score Regression (LDSC)^42^ to the meta-analyzed summary statistics to estimate the heritability and potential population stratification. We also performed cross-trait LDSC^43^ to compare the statistics from the LOGO GWAS (meta-analysis of 9 subgroup GWASs) and those from the conventional GWAS (using all the individuals in the cohort). The summary results of the GWASs are described in **Supplementary Table 4**.

#### UK Biobank

We applied the ten-fold LOGO approach to 20 clinical phenotypes for which individual-level phenotype data in UK Biobank was available (**Supplementary Table 7**). We performed GWASs using the linear regression model in Hail v0.2 (see **URLs**) with covariates including age, age^2^, sex, and the top 20 principal components. For blood pressure traits, we added 15 mmHg and 10 mmHg to sBP or dBP, respectively, if individuals are taking anti-hypertensive medication and derived the MAP and PP using the adjusted sBP and dBP. We also added smoking status as a covariate for blood pressure-related traits. We again performed cross-trait LDSC^43^ to compare the statistics from the LOGO GWAS and those from the conventional GWAS, for which we used summary statistics from Dr. Benjamin Neale’s lab (see **URLs**). The summary results of the meta-analyzed GWASs are described in **Supplementary Table 8**. For the additional 13 traits among the remainder of the 25 traits investigated in BioBank Japan, we were able to collect independent large-scale GWAS summary statistics of European ancestry, either from publicly available websites or upon request to the authors. The information of these 13 GWASs is described in **Supplementary Table 6**.

#### FinnGen

We did not perform within-cohort GWASs for the FinnGen cohort because the availability of individual-level phenotype data was limited. For the 20 traits where we performed LOGO in UK Biobank, we referred to UK Biobank GWAS summary statistics from all 361,194 white British individuals. With the exception of C-reactive protein (CRP), for 12 traits among the 13 traits where we used independent GWAS summary statistics in UK Biobank, we utilized the same GWAS summary statistics, as we confirmed that there was no apparent cohort overlap with FinnGen (**Supplementary Table 6)**. For CRP, since the GWAS of Ligthart et al. included the FINRISK Study, which was also involved in FinnGen, we additionally performed GWAS in UK Biobank individuals (*n* = 353,466). When performing CRP GWAS in UK Biobank, we excluded the individuals with autoimmune or inflammatory diseases.

### Construction of Polygenic Risk Scores

#### BioBank Japan

By referring to the effect sizes and *P* values of ten summary results from meta-analyzed GWASs of nine sub-group GWASs, we derived the PRSs of individuals in the one withheld sub-group using a clumping and thresholding method. First, we performed LD clumping on the meta-analyzed GWAS summary statistics with PLINK software using 5,000 randomly selected BioBank Japan participants as the LD reference. Briefly, we first used PLINK to clump all the variants using the following flags: --clump-p1 1 --clump-p2 1 --clump-r2 0.1 --clump-kb 1000. We then computed PRSs for variants meeting the following *P* value thresholds: 5×10^-8^, 5×10^-7^, 1×10^-6^, 1×10^-4^, 1×10^-3^, 1×10^-2^, 5×10^-2^, 0.1, 0.2, 0.5, and 1. In the one withheld sub-group, we derived PRSs by multiplying the dosage of risk alleles for each variant by the effect size in the GWAS and summing the scores across all the selected variants. We quantified the trait variance explained by the derived PRSs in individuals within the withheld sub-group, by calculating the adjusted R^2^ attributable to the PRSs from nested models, in which the full linear model was the trait value ∼ PRS + all covariates and the nested model dropped only the PRS term (**Supplementary Table 5**).

#### UK Biobank and FinnGen

For the clinical phenotypes for which the individual clinical data was available (20 traits in UK Biobank), we derived the PRSs in the same manner as described above for BioBank Japan (the ten-fold LOGO approach and deriving the PRSs in the one withheld group using the weights from the meta-analyzed summary statistics of nine sub-group GWASs by a clumping and thresholding approach). The variance explained by the derived PRSs is described in **Supplementary Table 10**. For the remaining 13 traits, we used a clumping and thresholding method on the collected large-scale GWAS summary statistics. Then, we derived the PRSs in the entire cohort referring to the weights and selected variants from the clumping and thresholding results. As noted above, we basically followed the original QC policy that had been adopted within each of the cohorts, and thus PRSs of UK Biobank and FinnGen could have included the rarer variants when compared with those of BioBank Japan (MAF > 0.0001 vs. MAF ≥ 0.01). We confirmed that both the performance of the PRSs and the result of downstream analyses did not substantially change, even when we restricted the variants used for calculating the PRSs to those with MAF ≥ 0.01 (i.e. the correlation *r* of these statistics exceeded 0.97) in UK Biobank and FinnGen.

### Survival Analysis using PRSs

#### BioBank Japan

We used Cox proportional-hazard models to test the association of the derived PRSs of clinical phenotypes with the length of lifespan (age at death) in the withheld sub-group. For the within-BioBank Japan analysis, we selected PRSs from the *P* value threshold of the best predictive capacity that had the largest variance explained by the PRS. We note that the threshold selection was based on the predictive capacity of the trait under investigation and not based on the result of the association of PRSs with lifespan. For the trans-biobank analysis, since there were no individual-level data available for some of the traits, optimization of the *P* value thresholds for all the traits was technically challenging. We thus selected PRSs from the *P* value threshold of 1.0×10^-6^, which was supposed to account for the polygenic architecture of complex traits while avoiding potential biases in PRS predictions induced by the large number of non-significant variants^44^. The PRSs for each trait in each sub-group were scaled to have zero mean and unit variance by Z-score transformation so as to obtain and compare the effect sizes across the investigated phenotypes. We used Cox proportional-hazard models to test the association of the scaled PRS of each trait in each sub-group with lifespan, with adjustment for sex, the 47-disease status and the top 20 principal components. We performed Schoenfeld residual tests^45^ to examine the proportional hazards assumption for the Cox regression. No apparent correlation between the Schoenfeld residuals and time was statistically and visually confirmed. We further meta-analyzed the association statistics from each of the ten sub-groups by the inverse variance method. A sex-stratified association study (**Supplementary Figure 4a**) was conducted by using the same Cox proportional-hazard models within male and female participants, except that we excluded sex from covariates.

To describe a standardized survival curve, we compared HRs for participants at the highest genetic risk (fifth quintile of PRSs) with those at an intermediate risk (quintiles 2 to 4) or the lowest risk (first quintile) as described previously^46^, which were standardized to the mean of all the covariates (**Supplementary Figure 7**). For the PRS of systolic blood pressure (sBP), we also analyzed the interaction effects with lifestyle factors recorded in the cohort. The lifestyle factors were obtained from the questionnaire to the participants, which asked them about their usual frequency of consumption or exercise of an investigated trait by selecting one from four categorical values. The answered values were converted to the quantitative values so that they represented the mean value of each category, except for the two binary lifestyle traits (whether the participant has ever smoked cigarettes and whether the participant currently drinks alcohol) (**Supplementary Table 12**). All the survival analyses were performed using the survival package in R software, version 3.3.0 (see **URLs**).

#### UK Biobank and FinnGen

For the quantitative traits where the individual level-data was available (20 traits in UK Biobank), we performed the same 10-fold survival analyses followed by meta-analysis as explained above in BioBank Japan. For the remaining traits, we performed the survival analyses on the entire cohort to test the association of the public GWAS-based PRS of each trait with lifespan. As described above, we adopted the *P* value threshold of 1×10^-6^ for the derivation of PRSs for the cross-biobank comparison. We included the same covariates used in the GWASs for each cohort, except for age and age^2^, in the Cox proportional hazard models. A sex-stratified association study (**Supplementary Figure 4b** and **c**) was conducted by using the same Cox proportional-hazard models within male and female participants, except that we excluded sex from the covariates.

As a secondary analysis, we performed a replication study of the association of sBP PRS on lifespan by using parental lifespan data in UK Biobank to validate the result of primary analysis with larger statistical power. To perform an association test of individuals’ genotype with their father’s and mother’s survival, we separately calculated Martingale residuals of the Cox model under a null model, scaled up to give a residual trait with a 1:1 correspondence with the HR, and tested its association with genotype dosage as described previously^47^.

For the PRS of BMI, we also analyzed the interaction effects with lifestyle factors recorded in UK Biobank. We collected the individual-level data of smoking status (ever smoked and smoking cessation), alcohol intake, coffee intake, and regular physical activity, and tested the effect of the interaction term between the BMI PRS and each of the lifestyle factors on lifespan.

We finally performed a fixed-effect meta-analysis of the PRS-lifespan association studies from BioBank Japan, UK Biobank, and FinnGen, by inverse variance method. To estimate the years of life gained or lost from PRS-lifespan associations, we converted the effect size from the Cox proportional hazard models into the years gained based on the following equation as described preciously^39, 47^;

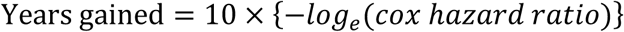

The association results of the trans-ethnic PRS meta-analysis including the years of life gained/lost are described them in **Supplementary Table 9**.

### Trans-ethnic Mendelian Randomization study

We conducted two-sample Mendelian randomization (MR) study to see the effect of each of 33 biomarkers on the outcome (i.e. lifespan) across three cohorts.

For the traits where we performed LOGO in PRS calculation (i.e. 33 traits in BioBank Japan and 20 traits in UK Biobank), we randomly split the cohort into half, and assigned them to the GWAS group (discovery) and the MR group (validation). For the selection of variants to be used as instrumental variables, we performed GWASs within the GWAS group for these traits with the same covariates described earlier, and selected independent genetic variants with *P*_GWAS_ < 1.0×10^-6^ for each trait (lead variants at significant loci at least +- 500 kb distant from each other). We next performed association study of these genetic variants with lifespan within the MR group, by using the same Cox proportional-hazard model described earlier. By using these genetic variants and association estimates, we obtained the effect estimate of the exposure (biomarker) on the outcome (lifespan) by pooling all MR estimates using the fixed-effects inverse-variance weighted method^48^.

For the traits where we used independent GWAS summary statistics in PRS calculation (i.e. 13 traits in UK Biobank and 33 traits in FinnGen), we selected independent genetic variants with *P*_GWAS_ < 1.0×10^-6^ from these statistics. We next performed association study of these genetic variants with lifespan in a whole cohort, by using the same Cox proportional-hazard model. These estimates are used to obtain the MR effect estimate by inverse-variance weighted method.

We finally performed the fixed-effect meta-analysis of these effect estimate in MR from each of the three cohorts.

## URLs

- Mean body mass index trends among adults estimates by country from Global Health Observatory data repository by WHO; http://apps.who.int/gho/data/view.main.BMIMEANADULTCv?lang=en
- UK Biobank; http://www.ukbiobank.ac.uk
- FinnGen; https://www.finngen.fi
- Source code for selecting individuals of white British ancestry; https://github.com/Nealelab/UK_Biobank_GWAS/blob/master/ukb31063_eur_selection.R
- Beagle4.1: https://faculty.washington.edu/browning/beagle/b4_1.html
- Hail software; https://hail.is
- Survival package in R software; https://cran.r-project.org/package=survival
- UK Biobank GWAS results from Dr. Benjamin Neale’s lab; http://www.nealelab.is/uk-biobank

